# Comprehensive gene expression analysis detects global reduction of proteasome subunits in schizophrenia

**DOI:** 10.1101/853226

**Authors:** Libi Hertzberg, Nicola Maggio, Inna Muler, Assif Yitzhaky, Michael Majer, Vahram Haroutunian, Pavel Katsel, Eytan Domany, Mark Weiser

**Affiliations:** Department of Physics of Complex Systems, Weizmann Institute of Science, Rehovot, Israel; Shalvata Mental Health Center, Affiliated to the Sackler School of Medicine, Tel Aviv University, Israel; Department of Neurology, The Chaim Sheba Medical Center, Tel Hashomer, Ramat Gan, Israel; Department of Neurology and Neurosurgery, Sackler Faculty of Medicine and Sagol School of Neuroscience, Tel Aviv University, Israel; Childhood Leukemia Research Institute and the Department of Pediatric Hemato-Oncology, Sheba Medical Center, Tel Hashomer, Ramat Gan, Israel; Human Molecular Genetics and Biochemistry, Faculty of Medicine, Tel Aviv University, Tel Aviv, Israel; Department of Psychiatry, Chaim Sheba Medical Center, Ramat-Gan and the Sackler School of Medicine, Tel-Aviv University, Israel; Departments of Psychiatry and Neuroscience, The Mount Sinai School of Medicine, New York, NY, USA; Department of Psychiatry, James J Peters VA Medical Center, Bronx, NY, USA

**Author notes:** **Correspondence:** Dr. Libi Hertzberg, Shalvata Mental Health Center, Affiliated to the Sackler School of Medicine, Tel Aviv University, Israel. 13 Aliat Hanoar St. Hod Hasharon 45100, Israel.

## Abstract

**OBJECTIVE:** A main challenge in the study of schizophrenia is its high heterogeneity. While it is generally accepted that there exist several biological mechanisms that may define distinct schizophrenia subtypes, they haven’t been identified yet. We applied comprehensive gene expression analysis, searching for molecular signals that differentiate patients with schizophrenia from healthy controls, and examined whether the identified signal characterizes a particular subgroup of the patients.

**METHODS:** We performed transcriptome sequencing of 14 superior temporal gyrus (STG) samples of relatively young (mean age: 44) subjects with schizophrenia and 15 matched controls from the Stanley Medical Research Institute. Analyses of differential expression and pathway enrichment were applied and the results were compared with those obtained from an independent cohort of elderly (mean age: 74) patients. Replicability was then tested on six additional independent datasets of various brain regions.

**RESULTS:** The two STG cohorts of relatively young and elderly subjects showed high replicability. Pathway enrichment analysis of the down-regulated genes pointed to proteasome-related pathways. Meta-analysis of differential expression identified down-regulation of 12 of 39 proteasome subunits in schizophrenia. Down-regulation of multiple proteasome subunits was replicated in six additional datasets (overall 8 cohorts, with 267 schizophrenia and 266 control samples, from 5 brain regions, were studied). This signal was concentrated in a subgroup of the patients.

**CONCLUSIONS:** We detect global down-regulation of proteasome subunits in a subgroup of the patients with schizophrenia. The proteasome is a major intracellular protein degradation system, where ubiquitinated proteins (proteins bound by the small protein called ubiquitin) are targeted for degradation. We hypothesize that the down-regulation we detect leads to proteasome dysfunction that causes accumulation of ubiquitinated proteins. Such accumulation has recently been identified, also in a subgroup of the studied patients with schizophrenia. Thus, down-regulation of proteasome subunits might define a biological subtype of schizophrenia.

## INTRUDOCTION

Schizophrenia affects 1% of the population and has a complex pathophysiology that is far from being fully understood. A main challenge in the study of schizophrenia is its high genetic and clinical heterogeneity (1). While for years several subtypes definitions were in scientific and clinical use, the DSM-5 has omitted them after concluding that they do not predict the course of illness (2). However, it is generally accepted that there exist several mechanisms that may define distinct schizophrenia subtypes, which haven’t been identified yet.

Recently, the ubiquitin mediated proteasome system (UPS), a protein degradation system, has been associated with schizophrenia in the transcript (3–6) and protein levels (7,8), with a down-regulation tendency in schizophrenia brain samples. In the genomic level, UPS pathways were enriched with schizophrenia associated copy number variants (9), and the proteasome pathway was enriched in schizophrenia susceptibility genes (10).

Recent findings suggest a more pronounced role of the UPS in schizophrenia. Accumulation of ubiquitinated proteins has been identified in brain samples of a subgroup of individuals with schizophrenia in the STG, frontal cortex and prefrontal cortex samples (11). Another recent study detected elevated ubiquitinated proteins levels in the orbitofrontal cortex of patients with schizophrenia (12). While ubiquitin binds to proteins (which are then “ubiquitinated”) to target them for proteasome degradation, proteasome dysfunction can cause accumulation of ubiquitinated proteins (13), as has been detected in schizophrenia. However, recent studies that have explored proteasome activity in schizophrenia have yielded inconsistent results (12,14). Thus, while elevation of ubiquitinated protein levels seems to play a role in schizophrenia, it is not clear whether this is caused by dysfunction of the proteasome.

Two studies (7,14) have examined protein levels of proteasome subunits in schizophrenia, where three regulatory subunits found to be decreased in both (see Table 2). Several studies reported down-regulation of proteasome subunits genes (4,6,15,16), but only two subunits were found to be down-regulated in more than a single study (see Table 2). Thus, while there is evidence for down-regulation of both transcript and protein levels of proteasome subunits in schizophrenia, the results are currently sporadic.

A basic limitation of gene expression studies of schizophrenia is the fact that brain samples are usually composed of a mixture of cell types, which might dilute authentic changes. In addition, as schizophrenia is highly heterogeneous (1), there are typically modest changes in gene expression (for example, (17)) which are thus difficult to detect. A relatively simple way to deal with these limitations is to perform a systematic comparison between independent datasets. Here we performed STG samples RNA-sequencing of 14 schizophrenia and 15 control subjects from the Stanley Medical Research Institute (SMRI). We applied pathway enrichment analysis to the list of genes detected as differentially expressed. We then used an independent cohort from the Mount Sinai School of Medicine (MSSM) to test the replicability of our results. A systematic meta-analysis of the SMRI and MSSM was applied to a subgroup of 39 inter-connected genes, which showed a tendency for down-regulation in schizophrenia. Six additional cohorts of different brain regions were used to further examine the robustness of our results. Note that one of the six datasets was from the same patients as the SMRI data described above, from a different brain region. Finally, we checked whether the signal characterizes a subgroup of the patients.

## METHODS

### Stanley Medical Research Institute (SMRI) subjects

STG postmortem tissues from 15 subjects with schizophrenia and 15 healthy controls were obtained from the SMRI using approved protocols for tissue collection and informed consent (18). Samples were examined by a neuropathologist to exclude cerebral pathologies (19). Diagnoses were performed independently by two psychiatrists according to DSM-IV, and matched by age, gender, post-mortem interval (PMI) and pH (Table 1). RNA-sequencing was applied to 29 out of the 30 STG samples (one sample did not pass quality control – see below).

**Table 1.** Subjects’ characteristics. Average values (standard deviation)

| Characteristics | Schizophrenia | Control |
| --- | --- | --- |
| **SMRI subjects** |  |  |
| Number of subjects | 14 | 15 |
| Gender (M/F) | 9/5 | 9/6 |
| Age (years) | 43.6 (13) | 48.1 (10.6) |
| Brain pH | 6.2 (0.3) | 6.3 (0.2) |
| **MSSM subjects** |  |  |
| PMI (minutes) | 2052 (900) | 1424 (596) |
| Number of subjects | 19 | 14 |
| Gender (M/F) | 14/5 | 5/9 |
| Age (years) | 77.4 (10.9) | 82.4 (12.7) |
| Brain pH | 6.4 (0.2) | 6.6 (0.3) |
| PMI (minutes) | 814 (499) | 460 (429) |

### MSSM subjects

STG samples of 19 schizophrenia and 14 healthy controls were obtained from the Brain Bank of the Department of Psychiatry of the MSSM (Table 1). All cortical dissections and sample preparation were described previously (20–22), see also the Supplementary information. Gene expression was measured using Affymetrix HG-U133A microarrays.

### RNA-sequencing

Brain regions were dissected at SMRI and delivered to Israel, where total RNA was isolated using the Trizol method. The concentration of total RNA and RNA Integrity Number (RIN) were measured. Samples with concentration ⩾ 10 ng/μl and RIN ⩾5 were selected for sequencing (one schizophrenia sample was excluded). Among these samples, the mean RIN was 6.3 (± 0.5), and the mean ratio of 260/280 was 1.6 (± 0.14). The mean total RNA yield was 15.4 μg (± 9.7). See Supplementary methods for a description of the libraries preparation protocol. For raw RNA-sequencing data description see Table 1S.

### Mapping, quantification of gene expression levels and pre-processing

We used standard software tools for mapping fragments to the genome and for quantification of gene expression levels. See supplementary methods for full description. Pre-processing: Lowess correction was calculated (23). Then expression threshold was set to 6 (log scale) to reduce noise. Filtering: Genes with expression values below 6 in at least 80% of the samples were filtered out of the analysis. 16,482 genes were left for the rest of the analysis after filtering (out of 23,715).

### MSSM microarray pre-processing

MAS-5 algorithm was used for normalization. Lowess correction was then applied, expression levels below 20 were set to 20 and log2-transformation was applied. Probe-sets without assigned gene symbols were removed. 12,033 probe-sets were left for the rest of the analysis after filtering (out of 22,283), representing 8,542 gene symbols. Probe sets of the same gene were combined. For full details see supplementary methods.

### Differential gene expression analysis

A linear model was fitted to each gene by a stepwise procedure (24), using the MATLAB function stepwiselm with default parameters. As pH did not differ significantly between schizophrenia and controls (Table 1), age, gender and PMI were included as covariates. The model was then refitted using only the selected variables, including diagnosis. Finally, for each gene, the diagnosis coefficient was statistically tested for being nonzero, implying an effect for schizophrenia, beyond any other effect of the covariates. This produced a t-statistic and a corresponding P-value. which were adjusted for multiple hypothesis testing using the false discovery rate (FDR) procedure (25). As the differentially expressed genes are subjected to further pathway enrichment analysis, a non-stringent FDR threshold of 15% was used. A standard 2-sample t-test was also performed; the results were very similar (Figure 1S).

**Figure 1.**
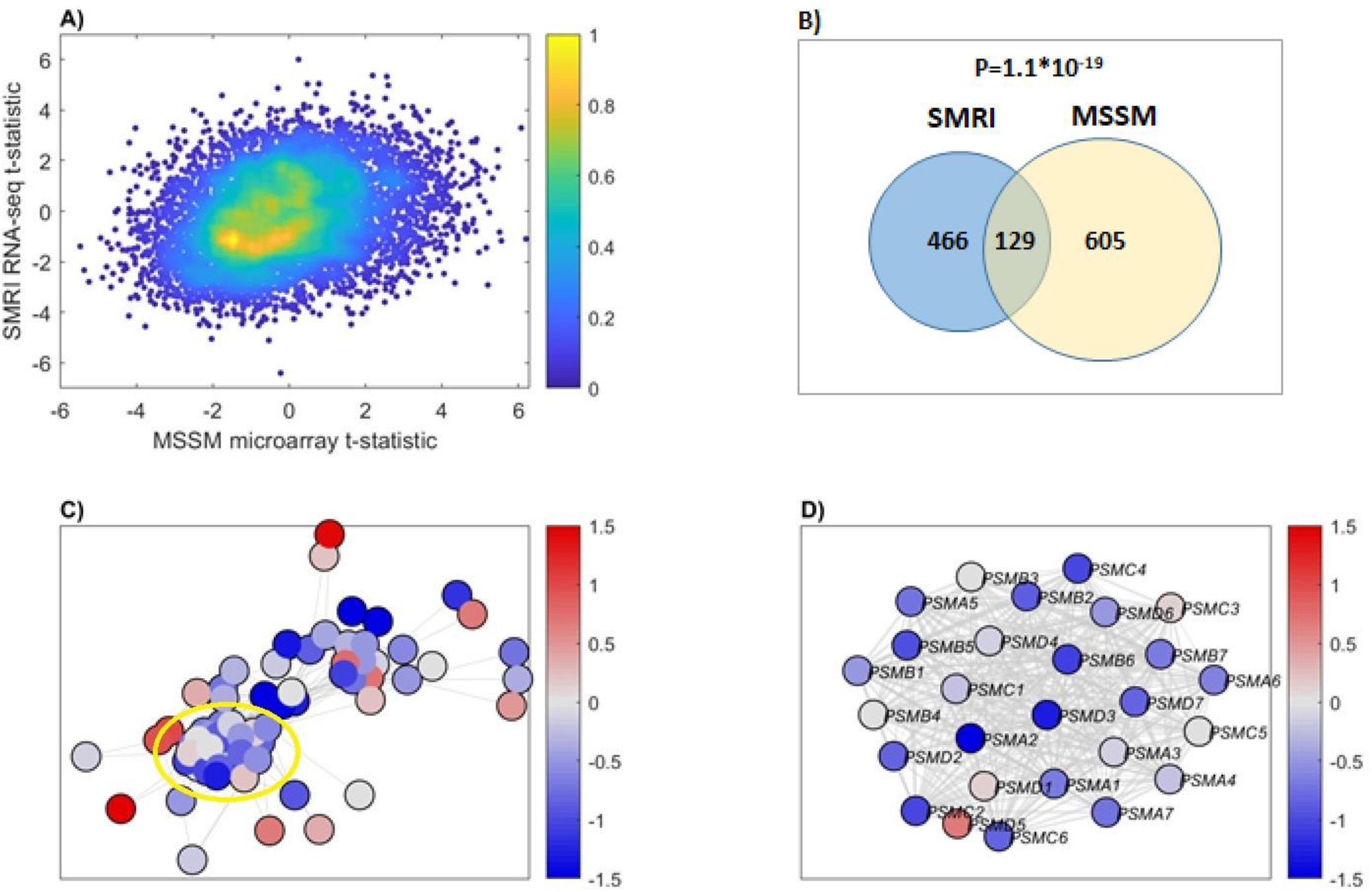
**A) Binned density scatter plot comparing the t-statistics for case versus control differential expression between the independent MSSM replication cohort assayed on microarrays and the SMRI RNA-seq data;** correlation between the statistics is 0.28 (P = 4.7*10-133). The colorbar represents the density in each cell, calculated by voronoi procedure (28) and normalized to values between 0 (minimal density) and 1 (maximal density). **B) Hypergeometric p-value calculation for the intersection between SMRI and MSSM down-regulated genes.** The 986 SMRI and 794 MSSM down-regulated genes were intersected with the 7,498 genes that are present in both cohorts, yielding 595 SMRI and 734 MSSM down-regulated genes, with 129 shared genes. **C) SMRI Differential expression network view for Ubiquitin-Proteasome Dependent Proteolysis superPathway.** A node’s color corresponds to the deviation of expression from the control samples group, in terms of standard deviation units. The edges represent STRING database co-expression relations. Only genes that have co-expression relations with other genes in the network are displayed. A subgroup of highly-interconnected genes, coding for proteasome subunits, is circled **D) Zoom in on proteasome subunits.** The same plot as in C), for a subgroup of highly-interconnected genes coding for proteasome subunits (circled in C))

### Pathway enrichment analysis using GeneAnalytics

GeneAnalytics tool (26) was used for pathway enrichment analysis. GeneAnalytics leverages PathCards (http://pathcards.genecards.org/), which clusters thousands of pathways from multiple sources into Superpathways, in order to improve inferences and reduce redundancy. Superpathways are scored by log2-transformation of the binomial p-value, which is equivalent to a corrected p-value with significance level <0.05.

### Differential expression STRING database network view

#### Network creation

Given a list of genes, a network is built. A network consists of genes (nodes) and genes’ co-expression relations (edges). The co-expression relations data was downloaded from the STRING database, version 10.5 (27). Each such connection has a score between 0 and 1 that “indicates the estimated likelihood that a given interaction is “biologically meaningful, specific and reproducible” (27). The product of this process is a network whose nodes correspond to genes and edges corresponds to co-expression relations. Only edges with STRING score greater than 0.1 are considered.

#### Differential expression network view

Given a network and gene expression data, of both patients and controls, the following steps are taken, for each gene:

1. The mean expression and standard deviation values, Mc and Sc, are calculated using the control samples only.
2. The mean expression, Mp, is calculated using the patients’ samples.
3. Mp-Mc is calculated, the difference in the expression means between the two groups of samples.
4. The deviation from the control group is calculated, by: (Mp-Mc)/Sc

Finally, the network is displayed as an undirected graph, with each node colored according to the deviation described above, (Mp-Mc)/Sc. The edges represent co-expression relations. Only genes that have co-expression relations with other genes in the network are displayed.

## RESULTS

### UPS related pathways are enriched in the group of genes which are down-regulated in SMRI STG samples of individuals with schizophrenia

Differential expression analysis was performed, yielding 881 up-regulated and 986 down-regulated genes. In order to examine possible connection to antipsychotic medications, alcohol or substance use, we performed correlation analyses between the expression pattern of the differentially expressed genes and Fluphenazine equivalent dosage, substance use and alcohol use measures. Correlation analyses for Fluphenazine equivalent dosage and alcohol use did not reveal any significant association with differential expression. Correlation analysis for substance use detected two down-regulated genes (out of 986) with statistically significant correlated expression (supplementary methods and Figures 2S-4S).

**Figure 2.**
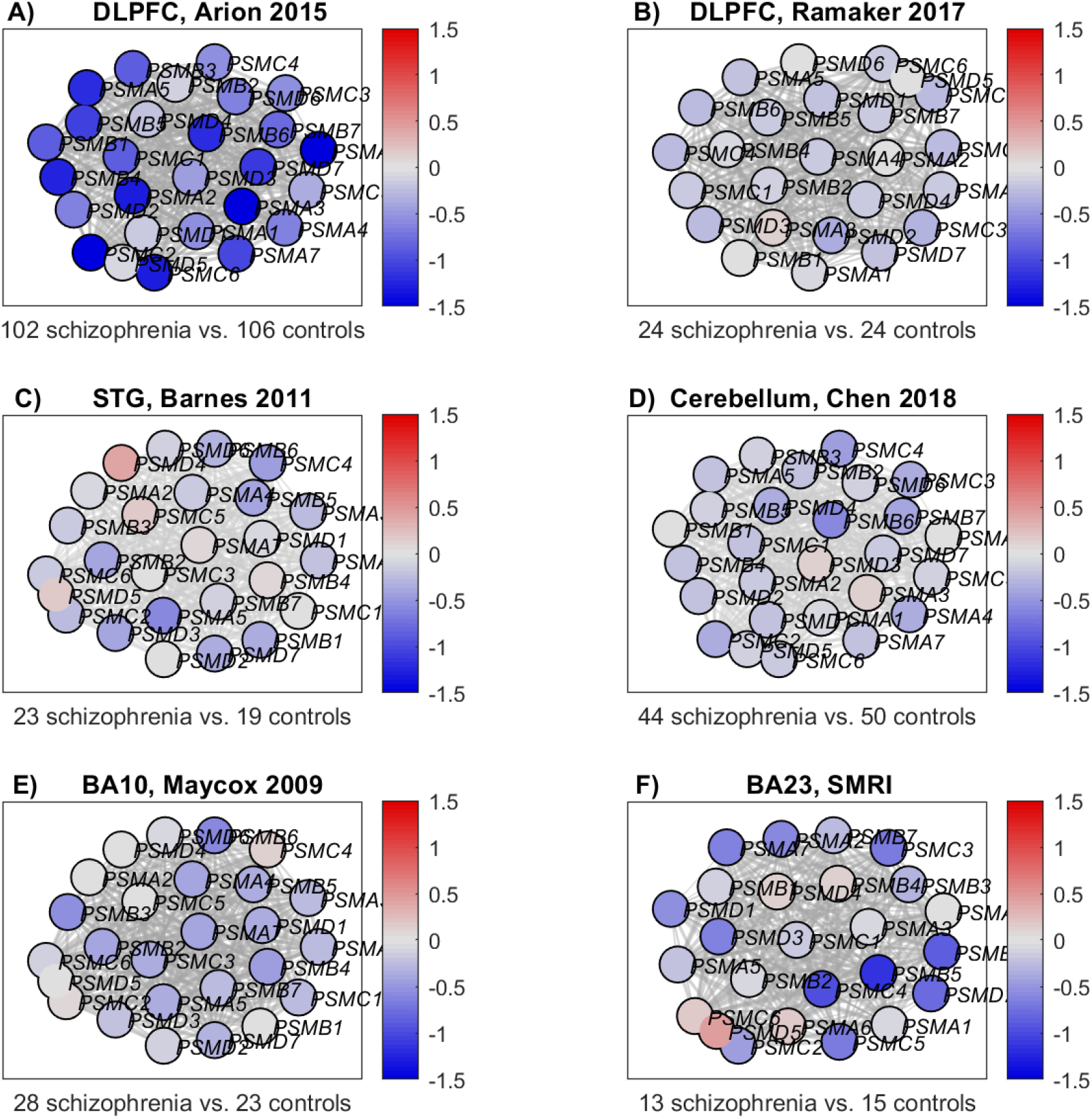
Proteasome subunits differential expression network view: The nodes’ colors correspond to the deviation from the group of the control samples, in terms of standard deviation units. The edges represent STRING database co-expression relations. Only genes that have co-expression relations with other genes in the network are displayed. **A) DLPFC, Arion 2015 dataset** (6). **B) DLPFC, Ramaker 2017 dataset** (31). **C) STG, Barnes 2011 dataset** (32). **D) Cerebellum, Chen 2018** (33). **E) BA10, Mycox 2009 dataset** (34) **F) BA23, SMRI dataset**

Pathway enrichment analysis was applied separately to the up and down-regulated genes. Results are presented in Tables 3S-4S for the up-regulated and down-regulated genes, respectively. Out of 49 pathways enriched in the down-regulated genes, five are directly UPS related (marked in Table 4S). While several pathways have higher enrichment scores, we focus on the UPS and proteasome-related pathways, since five such pathways were enriched, and several closely related additional pathways were also enriched (see Table 4S).

### The UPS signal is highly replicated in the MSSM STG samples

We tested whether our findings are replicated in the STG of the MSSM cohort, an independent cohort of elderly subjects. We first examined whether the two datasets are comparable. Though microarrays differ from RNA-seq in their capture features, there was a significant positive correlation of the t-statistics (schizophrenia vs. controls) between SMRI and MSSM across 7,498 genes common to both platforms (Figure 1A).

We next repeated the differential expression and pathway enrichment analyses in the MSSM cohort. 919 genes and 794 genes were found to be up-regulated and down-regulated in schizophrenia, respectively. MSSM and SMRI differentially expressed genes significantly overlap (hypergeometric P-values: 9.8*10^−7^, 1.1*10^−19^ for the up-regulated and down-regulated genes, respectively; see Figure 1B).

Pathway enrichment analysis was applied, and 27 and 48 pathways were enriched in up-regulated and down-regulated genes, respectively (results are listed in Tables 5S and 6S). Intersection between SMRI 49 enriched pathways and MSSM 48 enriched pathways in the down-regulated genes yields 30 pathways, listed in Table 4S (hypergeometric p-value: 2.5*10^−36^). Four out of the five SMRI enriched UPS pathways were enriched also in the MSSM. A similar analysis of the up-regulated genes yields a hypergeometric p-value of 1.03*10^−6^.

### A network view of the UPS identifies down-regulation of a tightly connected cluster of proteasome subunits

To further explore the UPS differential expression, we applied differential expression network view to the SMRI (see methods). It was applied to the Ubiquitin-Proteasome Dependent Proteolysis GeneAnalytics “superpathway” (26), which is representative of the UPS and was significantly enriched in both SMRI and MSSM (see Table 4S). The network view includes all 69 pathway genes for which network data was available from STRING (29), and not only those 27 genes that were found to be down-regulated. As can be seen in Figure 1C, there is a cluster of tightly inter-connected genes which are mostly down-regulated in schizophrenia (bluish colours of the nodes). Interestingly, this cluster is composed of proteasome subunits, as shown in Figure 1D. The same analysis of the MSSM yields a similar view (Figure 5S).

### Meta-analysis of SMRI and MSSM datasets identifies down-regulation of multiple proteasome subunits in STG samples of subjects with schizophrenia

We performed a meta-analysis of the expression of each of the 39 proteasome subunit genes, whose expression has been measured by both SMRI and MSSM (see supplementary methods). The list of proteasome subunits genes, meta-analysis results and comparisons to previous gene expression and protein-level studies are summarized in Table 2. Overall 12 out of 39 subunit genes were found to be down-regulated.

**Table 2.** Proteasome subunits differential gene and protein level expression, in previous studies and in our meta-analysis. Previous gene expression studies’ results were listed only for genes which were detected as differentially expressed in more than one study. Down-regulation findings are highlighted in blue. In the meta-analysis, a gene is defined as down-regulated if its summary measure is lower than zero and the confidence interval doesn’t cross zero

| # | Proteasome subunit genes | Previous gene expression studies | Previous protein level studies (7,14) | Our meta-analysis (SMRI+MSSM) | SMRI+ MSSM meta-analysis summary measure [confidence interval] |
| --- | --- | --- | --- | --- | --- |
| **Structural subunits** |  |  |  |  |  |
| **20S core $\alpha$ subunits** |  |  |  |  |  |
| 1 | PSMA1 (also named 20S α1) | Down-regulated in 2 studies (4,30) | Not measured | unchanged | -0.37 [-0.97, 0.21] |
| 2 | PSMA2 (20S α2) | | Not measured | Down-regulation | -1.13 [-1.68, 0.59] |
| 3 | PSMA3 (20S α3) | | Not measured | unchanged | -0.63 [-1.67, 0.4] |
| 4 | PSMA4 (20S α4) | | Not measured | unchanged | -0.43 [-0.9, 0.07] |
| 5 | PSMA5 (20S α5) | | Not measured | Down-regulation | -0.61 [-1.13, -0.09] |
| 6 | PSMA6 (20S α6) | | unchanged in (14) | Down-regulation | -0.63 [-1.15, -0.12] |
| 7 | PSMA7 (20S α7) | | Not measured | Down-regulation | -0.79 [-1.32, 0.27] |
| **Catalytic subunits** |  |  |  |  |  |
| **20S core $\beta$ subunits** |  |  |  |  |  |
| 8 | PSMB1 (20S β1) | | Not measured | unchanged | -0.17 [-0.73, 0.37] |
| 9 | PSMB2 (20S β2) | | Down-regulation trend (7) (p=0.08); unchanged in (14) | Down-regulation | -0.62 [-1.13, 0.11] |
| 10 | PSMB3 (20S β3) | | Not measured | unchanged | -0.25 [-0.75, 0.25] |
| 11 | PSMB4 (20S β4) | | Not measured | unchanged | -0.03 [-0.53, 0.46] |
| 12 | PSMB5 (20S β5) | | unchanged (7,14) | Down-regulation | -0.73 [-1.28, -0.18] |
| 13 | PSMB6 (20S β6) | | Not measured | unchanged | -0.13 [-1.25, 0.97] |
| 14 | PSMB7 (20S β7) | | Not measured | unchanged | -0.37 [-0.87, 0.13] |
| **Immunoproteasome $\beta$ subunit genes** |  |  |  |  |  |
| | PSMB8 (20S β5i) | | unchanged (7,14) | unchanged in | |
|  |  |  |  | SMRI; absent in MSSM |  |
| 15 | PSMB9 (20S β1i) | | unchanged (7) | unchanged | -0.04 [-0.45, 0.54] |
| 16 | PSMB10 (20S β2i) | | unchanged (7,14) | unchanged | 0.16 [-0.38, 0.71] |
|  | **Regulatory subunits** |  |  |  |  |
|  | **19S AAA-ATPase subunits (Rpt)** |  |  |  |  |
| 17 | PSMC1 (19S Rpt2) |  | unchanged in (7); **Down-regulated** in (14) | unchanged | 0.02 [-0.48, 0.52] |
| 18 | PSMC2 (19S Rpt1) |  | **Down-regulated** in two studies (7,14) | **Down-regulation** | -0.93 [-1.46, 0.4] |
| 29 | PSMC3 (19S Rpt5) |  | Unchanged in (7); **Down-regulated** in (14) | unchanged | -0.11 [-0.62, 0.38] |
| 20 | PSMC4 (19S Rpt3) |  | **Down-regulated** in two studies (7,14) | **Down-regulation** | -0.67 [-1.19, 0.15] |
| 21 | PSMC5 (19S Rpt6) |  | **Down-regulated** in two studies (7,14) | unchanged | 0.02 [-0.48, 0.52] |
| 22 | PSMC6 (19S Rpt4) | **Down-regulated** in two studies (4,6) | unchanged (7); **Down-regulated** in (14) | **Down-regulation** | -0.83 [-1.36, 0.3] |
|  | **19S non-ATPase subunits (Rpn)** |  |  |  |  |
| 23 | PSMD1 (19S Rpn2) |  | Not measured | unchanged | 0.07 [-0.42, 0.58] |
| 24 | PSMD2 (19S Rpn1) |  | Not measured | unchanged | -0.03 [-1.49, 1.41] |
| 25 | PSMD3 (19S Rpn3) |  | Not measured | unchanged | -0.34 [-0.87, 0.18] |
| 26 | PSMD4 (19S Rpn10) |  | unchanged (7) | unchanged | 0.09 [-0.41, 0.59] |
| 27 | PSMD5 |  | Not measured | unchanged | 0.17 [-0.69, 1.05] |
| 28 | PSMD6 (19S Rpn7) |  | Not measured | **Down-regulation** | -0.62 [-1.13, -0.1] |
| 29 | PSMD7 (19S Rpn8) |  | Not measured | unchanged | -0.07 [-1.02, 0.86] |
| 30 | PSMD8 (19S Rpn12) |  | Not measured | unchanged | -0.39 [-0.9, 0.11] |
| 31 | PSMD9 (19S Rpn4) |  | Not measured | unchanged | -0.39 [-1.42, 0.63] |
| 32 | PSMD10 |  | Not measured | unchanged | 0.13 [-0.36, 0.63] |
| 33 | PSMD11 (19S Rpn6) |  | unchanged (7) | **Down-regulation** | -0.73 [-1.25, -0.21] |
| 34 | PSMD12 (19S Rpn5) |  | Not measured | unchanged | -0.23 [-0.74, 0.26] |
| 35 | PSMD13 (19S Rpn9) |  | Not measured | unchanged | 0.05 [-0.58, 0.69] |
| 36 | PSMD14 (19S Rpn11) |  | unchanged (7) | **Down-regulation** | -0.89 [-1.42, -0.37] |
|  | **11S subunits** |  |  |  |  |
| 37 | PSME1 (11S $\alpha$) | | **Down-regulated** in (7); unchanged in (14) | unchanged | -0.33 [-0.84, 0.16] |
| 38 | PSME2 (11S $\beta$) | | unchanged (7,14) | unchanged | 0.29 [-0.65, 1.24] |
| 39 | PSME3 (11S gamma) |  | unchanged (7) | unchanged | -0.07 [-0.85, 0.7] |

### Down-regulation signal of proteasome subunits in schizophrenia is replicated in 6 independent datasets of 5 different brain regions

To examine whether the down-regulation of proteasome subunits is specific to the STG we repeated the differential expression network analysis of the 39 proteasome subunit genes using 6 additional datasets (fully described in the supplementary information): dorso-lateral prefrontal cortex (DLFPC) samples from Arion 2015 (6) and from Ramaker 2017 (31), STG samples from Barnes 2011 (32), Cerebellum samples from Chen 2018 (33), Brodmann area 23 (BA23) samples from the SMRI cohort, and Brodmann area 10 (BA10) samples from Mycox 2009 (34). The results are presented in Figure 2. The DLPFC samples of Arion 2015 (Figure 2A) exhibit pronounced down-regulation, while in the DLPFC samples of Ramaker 2017 (Figure 2B) the signal is weaker, though present in most of the genes; the binomial p-value for the number of genes with (even slightly) reduced expression versus the control group is p= 6.4 10^−6^). Interestingly, while Ramaker 2017 (31) used brain samples composed of mixture of cells, Arion 2015 (6) used laser microdissection to capture pyramidal neurons. Thus, the difference in the down-regulation magnitude might occur due to dilution of the signal, caused by the mixture of cell types used in Ramaker 2017. The Cerebellum samples from Chen 2018 (33) (Figure 2D), BA10 samples from Mycox 2009 (Figure 2E) and BA23 SMRI (Figure 2F) show clear tendency for down-regulation (binomial p-values 6.9 10^−7^, 1.2 10^−5^ and 0.04, respectively), with modest magnitude (mostly less than 1 standard deviation). STG samples of Barnes 2011 (32) (Figure 2C) show a similar pattern. Down-regulation might be specific to neurons or subtypes of neurons; as the brain samples in these datasets are of mixture of cells, the signal might be diluted. Overall, this analysis replicates the signal of down-regulation of multiple proteasome subunits, in both the STG and additional 4 brain regions.

### Down-regulation of proteasome subunits in schizophrenia is concentrated in a subgroup of the patients

To explore whether the signal is concentrated in a subgroup of the patients, we applied fold change analysis of the 12 down-regulated proteasome subunits (listed in Table 2) to each of the SMRI schizophrenia samples. The results are plotted in Figure 3A. Half of the patients (7/14, “Group 2”; marked blue along the x-axis) show down-regulation tendency (bluish colors) of most of the 12 proteasome subunits genes, while the others (“Group 1”; marked green) show fold change values closer to 1, for most genes. When applying the same analysis to the Arion 2015 dataset, of microdissected pyramidal neurons, an even more pronounced picture is seen (Figure 3B). A similar picture is seen for the other 6 datasets (Figure 7S). Support for this observation comes from a recent study (35), where transcriptomics analysis of 189 DLPFC samples of patients with schizophrenia vs. 206 healthy controls identified two molecular subtypes of schizophrenia. In “Type 1” (about half of the patients) four differentially expressed genes (schizophrenia vs. controls) were detected, and in “Type 2” more than 3000. When examining the list of differentially expressed genes (Supplemental Table 3B), 28 proteasome subunits were differentially expressed, all down-regulated, in “Type 2”, while no proteasome subunit genes were differentially expressed in “Type 1”.

**Figure 3.**
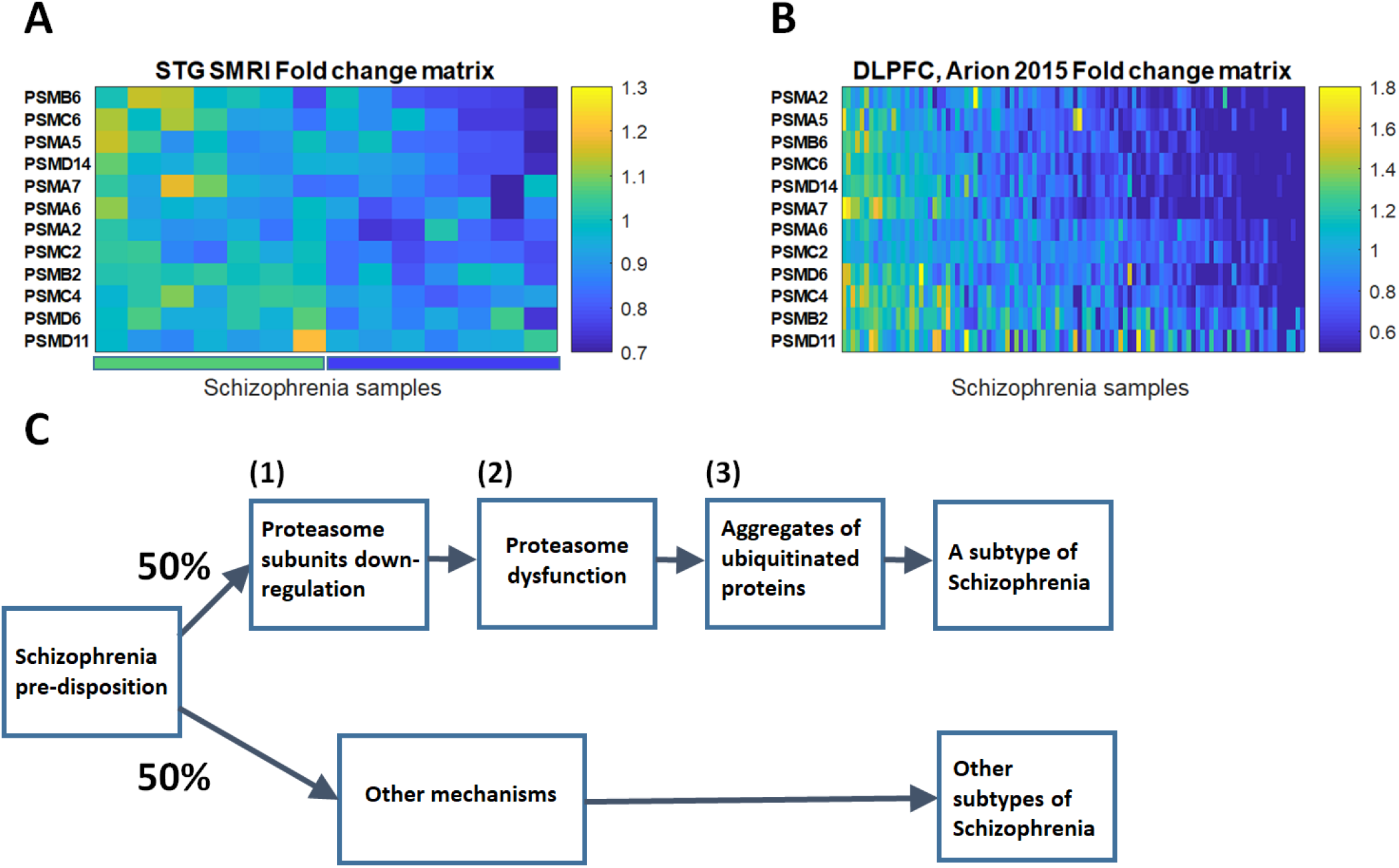
**A) SMRI STG schizophrenia samples fold change matrix of proteasome subunits genes.** Each row represents one of the 12 proteasome subunits genes that were found to be down-regulated in schizophrenia in the meta-analysis of the SMRI and MSSM datasets. Each column represents one of the SMRI schizophrenia samples. The color code represents the fold change, i.e. the expression value of the proteasome subunit gene in the specific sample, divided by its mean expression in the 15 control samples. Samples and genes locations were sorted by the SPIN tool (36). The left half of the samples, “Group 1”, are marked by a green bar along the x-axis and the right half, “Group 2”, with blue. **B) DLPFC Arion 2015 schizophrenia samples fold change matrix of proteasome subunits genes.** The same plot as in A) for the DLPFC Arion 2015 dataset. **C) A schematic preliminary model for a biological mechanism based division of schizophrenia patients into subtypes**.

We then applied a similar analysis as in (35), and compared each of “Group 1” and “Group 2” samples to the controls, separately. Differential expression analysis was applied to the 47 SMRI measured proteasome subunit genes, in each of the two groups. While in “Group 1” no differentially expressed genes were found, in “Group 2” 23 proteasome subunits were found to be differentially expressed (FDR < 15%; Table 8S). We conclude that the signal of proteasome subunits down-regulation characterizes about half of the patients with schizophrenia.

## DISCUSSION

The main finding of our study is a global down-regulation of multiple proteasome subunits in post mortem brain samples of individuals with schizophrenia. Although several scenarios may be possible, a reasonable model (Figure 3C) is that given a predisposition to schizophrenia, certain (unknown) factors lead to (1) down-regulation of multiple proteasome subunits in about half of the patients. This in turn leads to (2) proteasome dysfunction which causes (3) accumulation of ubiquitinated proteins. We discuss below the evidence relevant to each of the hypotheses (1)-(3) of our model.

Hypothesis (1) relates directly to our main finding, which was replicated in 8 datasets of 5 different brain regions. We observed that the signal characterizes about half of the patients. This observation is supported by (35), where two molecular subtypes of schizophrenia were detected, one (“Type 2”) with 28 down-regulated proteasome subunits genes, and one (“Type 1”) without dysregulation of proteasome subunits genes. It should be noted that in “Type 2” more than 3000 genes (about 25% of the measured genes) were found to be dysregulated. Thus, the fact that 28 proteasome subunits genes were found to be dysregulated is somewhat less surprising. However, as all the 28 were down-regulated, it makes the concordance with our results significant.

The fact that several studies identified decreased protein levels of proteasome subunit genes (Table 2) supports hypothesis (2), of proteasome dysfunction in schizophrenia. However, it wasn’t established whether the lower protein levels are caused by lower expression of the coding genes. Moreover, previous studies of proteasome activity in schizophrenia yielded inconsistent results. While in (14) intra-cellular compartment-specific dysfunction in STG samples was found, no change has been detected in neither blood or brain in (12). A possible explanation of this inconsistency is that the signal is specific not only to a subgroup of the patients, but also to neurons or subtypes of neurons, and thus diluted. This is supported by our analysis of Arion 2015 dataset, of laser microdissected neurons (6), where higher magnitude of down-regulation was detected (Figure 2). This could also explain why the signal hasn’t been detected by many previous relevant gene expression studies. Actually, if we look at some of the datasets (for example, in Figure 2B-E), each proteasome subunit is not pronouncedly down-regulated. Only the analysis of the proteasome subunits as a group, measured in multiple datasets, enabled the detection of the global down-regulation signal.

Hypothesis (3), of accumulation of ubiquitinated proteins in schizophrenia, comes from two recent studies. In (11), accumulation of ubiquitinated proteins has been identified in brain samples of about half of the patients in the STG, frontal cortex and prefrontal cortex samples. In (12), ubiquitinated protein levels were found to be elevated in the orbitofrontal cortex of patients with schizophrenia. While the fact that dysfunction of proteasome can cause accumulation of ubiquitinated proteins (13) suggests a causative connection between hypotheses (2) and (3), this link wasn’t examined in schizophrenia.

As described in (11), the accumulated ubiquitinated proteins were enriched with nervous system development related pathways, suggesting its possible relation to disease pathogenesis through disruption of relevant pathways. In addition, clinical symptoms were correlated with two ubiquitin conjugation genes’ expression in patients’ peripheral blood (37). These findings might suggest that our hypothesized model defines a biological and clinical subtype of schizophrenia. In this context we note that Bortezomib, a proteasome inhibitor used in the treatment of cancer, is not known to cause psychosis when given to glioblastoma patients (where the brain-blood-barrier is disrupted) (38–40). This seemingly suggests that proteasome dysfunction is not the cause of the symptoms seen in schizophrenia. However, additional studies are needed in order to further explore this connection.

Our study is limited by several features. Every postmortem study represents only a snapshot at the end of life. This is especially relevant in schizophrenia, as its pathogenesis is probably rooted in early development (41). The fact that we compare independent cohorts of both relatively young and elderly subjects strengthens the validity of the results, but doesn’t fully overcome this limitation. There is also the question of pharmacotherapy, as exposure to antipsychotics might affect gene expression. We found no significant correlation between Fluphenazine equivalent dose and expression levels. In addition, the fact that the subjects of the cohorts significantly differ in age suggests that duration of exposure to antipsychotics is unlikely to influence proteasome subunits expression substantively. The replication of the detected signal in 8 cohorts from 5 brain regions significantly increases the validity and generalizability of this signal. As gene expression does not always correlate with the levels of the coded proteins, the fact that we measure gene expression alone is a serious limitation, which causes difficulties in making definitive conclusions regarding the biological consequences of the results. While several studies detected decreased protein levels of proteasome subunits (7,14), the results were not fully consistent and the recent proteasome activity studies in schizophrenia were not consistent either, as described above. Thus, further study is needed in order to decipher the consequences of the global down-regulation of proteasome subunits we detect in schizophrenia, in terms of protein levels and proteasome activity.

Overall, we detect global down-regulation of proteasome subunits in schizophrenia, which characterizes about half of the patients. Based on ours and others’ recent findings we present a hypothesized model for a mechanism that defines a biological, and maybe also clinical, subtype of schizophrenia. This has the potential to lead to a better understanding of the biological and clinical subtypes of schizophrenia and to finding novel diagnostic and therapeutic tools.

