## Supplementary Information for "Comprehensive gene expression analysis detects global reduction of proteasome subunits in schizophrenia"

### LIST OF FIGURES

Figure 1S: Comparison between multiple linear regression and t-test analysis resulting SMRI STG up-regulated and down-regulated genes. A) Venn diagram for the intersection between the 881 genes that were found to be up-regulated in schizophrenia STG SMRI samples using multiple linear regression analysis (with FDR Q < 15%) and 855 genes that were found to be up-regulated in schizophrenia STG SMRI samples using t-test analysis (FDR Q < 15%). B) Venn diagram for the intersection between the 986 genes that were found to be down-regulated in schizophrenia STG SMRI samples using multiple linear regression analysis (with FDR Q < 15%) and 944 genes that were found to be down-regulated in schizophrenia STG SMRI samples using t-test analysis (FDR Q < 15%). .......... 10

Figure 5S: MSSM STG Differential expression network view: A) Ubiquitin-Proteasome Dependent Proteolysis superPathway. The node's colors correspond to the deviation from the control samples group, in terms of standard deviation units (see Methods). The edges represent STRING database co-expression relations. Only genes that have co-expression relations with other genes in the network are displayed. A subgroup of highly-interconnected genes, coding for proteasome subunits, is circled. B) Zoom in on proteasome subunits. The same plot as in A), for a subgroup of highly-interconnected genes coding for proteasome subunits (circled in A)). .......... 11

Figure 6S: PSMA5 STG SMRI + MSSM meta-analysis of differential expression. Forest plot was generated using the function "forest" from the "meta" package in R, version 4.9-2 (General Package for Meta-Analysis) (8). The forest plot shows the differences in PSMA5 expression between subjects with schizophrenia and healthy controls, for each of the two studies, SMRI and MSSM. Each square represents the standardized difference (Hedges' g (7)) between schizophrenia and control for that study, with the area of the square reflecting the weight (determined by the sample size) given to that study in the meta-analysis. Each horizontal line represents the 95% confidence interval for the mean difference in that study. The vertical line shows the point of 0 difference. The standardized difference is positive (negative) if the expression is higher (lower) in schizophrenia vs. the control group. The center of the diamond represents the overall difference across both studies and its width represents 95% confidence interval. ..... 12

Figure 7S: Fold change matrix of proteasome subunits genes. A) Cerebellum, Chen 2018 data (47). Each row represents one of the 12 proteasome subunits genes that were found to be down-regulated in schizophrenia in the meta-analysis of the SMRI and MSSM datasets. Each column represents one of the Chen 2018 44 schizophrenia samples. The color code represents the fold change, i.e. the expression value of the proteasome subunit gene in the specific sample, divided by its mean expression in the 50 control samples. B) DLPFC, Ramaker 2017 dataset (45). 24 samples of schizophrenia patients vs. 24 controls. C) DLPFC, Arion 2015 dataset (6). 102 samples of schizophrenia patients vs. 106 controls. D) BA10, Mycox 2009 dataset (48). 28 samples of schizophrenia patients vs. 23 controls. E) STG SMRI dataset. 14 samples of schizophrenia patients vs. 15 controls. F) BA23 SMRI dataset. 13 samples of schizophrenia patients vs. 15 controls. ..... 13

Table 4S. Pathway enrichment analysis of SMRI STG down-regulated genes with corrected p-value < 0.05. GeneAnalytics tool superpathways that were found to be enriched in the list of down-regulated genes are ordered by descending order of their enrichment score. The enrichment scores are in the second column and the superpathways' names are listed in the third column. The fourth column presents the number of down-regulated genes that belong to each superpathway , with the total number of genes of the superpathway in parentheses. MSSM enrichment score is given in the 5th column, where (-) sign means that the superpathway wasn't enriched in the list of MSSM down-regulated genes. For superpathways that are known to involve the UPS, a reference indicating the UPS involvement is given in the 6th column. Ubiquitin-proteasome directly related pathways are in bold ..... 15

Table 5S: Pathway enrichment analysis of MSSM STG up-regulated genes with corrected p-value < 0.05. GeneAnalytics tool superpathways that were found to be enriched in the list of up-regulated genes are ordered by descending order of their enrichment score. The number of up-regulated genes that belong to each superpathway is given, with the number of genes of each superpathway in parentheses. ..... 17

Table 6S: Pathway enrichment analysis of MSSM STG down-regulated genes with corrected p-value < 0.05. GeneAnalytics tool superpathways that were found to be enriched in the list of down-regulated genes are ordered by descending order of their enrichment score. The number of down-regulated genes that belong to each superpathway is given, with the number of genes of each superpathway in parentheses. UPS directly related pathways are in bold ..... 18

Table 7S: STG down-regulated genes enriched UPS-related pathways; numbers of SMRI and MSSM hits. For each of the 5 UPS-related pathways that were enriched in the SMRI STG down-regulated genes, the number of 'hits' (down-regulated genes that belong to the pathway) shared with MSSM STG down-regulated genes is given in the second column. The numbers of hits specific to SMRI and MSSM down-regulated genes are given in columns 3 and 4, respectively. 19

Table 8S: SMRI STG Differential expression analysis of proteasome subunits genes. 7 schizophrenia samples ("Group 2") were compared to the 15 control samples. T-sample t-test was applied; p-values were corrected by the benjamini-Hochberg procedure (6) ..... 19

#### *Mount Sinai School of Medicine (MSSM) subjects*

Human brain samples of 19 schizophrenia and 14 healthy controls of the superior temporal gyrus (STG) were obtained from the Brain Bank of the Department of Psychiatry of the MSSM (Table 1). All cortical dissections and sample preparation were described previously (1–3). Brain banking activities were approved by the MSSM Institutional Review Board and written consent for brain donation was obtained from the next-of-kin of all subjects. Cases diagnosed as schizophrenia met the DSM-III/IV criteria, as determined by clinical investigators. None of the samples, of neither subjects with schizophrenia nor controls, showed evidence of any significant neuropathology (4). Whole-genome gene expression was measured using Affymetrix HG-U133A microarrays.

#### *MSSM Superior Temporal Gyrus (STG) Gene expression preprocessing*

The HG-U133A Affymetrix chips were pre-processed with the commonly used Affymetrix MicroArray Suite v. 5.0 (MAS-5) algorithm. MAS-5 was criticized for its high false positive rate claimed to stem from making use of mismatches, as opposed to robust multi-array average. However, it was shown that combined with detection calls, MAS-5 is both selective and sensitive (5). Lowess correction was then calculated. As we observed a random-like distribution for probe-sets with low expression levels, we set all the expression levels below 20 to be 20. Data were subjected to log2-transformation. Filtering: Probe-sets that are present in at least 40% of the samples of at least one of the *17* regions of a certain disease type (schizophrenia or control) are kept for the rest of the analysis. Probe-sets without assigned Affymetrix gene symbols annotation were removed. 12,033 probe-sets were left for the rest of the analysis after filtering (out of 22,283), representing 8,542 gene symbols.

#### *MSSM STG Combining probe sets of the same gene*

Genes represented by more than one probe-set with the same Affymetrix assigned gene symbol were considered to represent the same gene and the expression was determined as follows: The Pearson correlation coefficient was calculated for each pair in such a group of probe-sets; and the largest subgroup in which each pair of probe-sets had a correlation coefficient higher than 0.5 was found by simple scanning. If the size of the chosen subgroup was larger than 2, the probe-set with the maximal average correlation values (in respect to the rest of the probe-sets in the subgroup) was chosen to represent the gene. Otherwise, in case the size of the chosen subgroup equals 2, one of them is chosen by random.

#### *Comparison between SMRI STG multiple linear regression and t-test analysis results*

In addition to multiple linear regression analysis, differentially expressed genes were identified by applying 2-sided t-test for each gene, comparing its expression between the schizophrenia samples and the controls. P-values were then adjusted for multiple hypothesis testing using false discovery rate (FDR) estimation (6), and the differentially expressed genes were determined as those with an estimated FDR ≤ 15%.

The comparison between the resulting lists of up-regulated and down-regulated genes, using multiple linear regression and t-test, is plotted in Figure 1S. It can be seen that the intersection between the lists is very large (calculated hyper-geometric p-value < $1*10^{-50}$, for both up-regulated and down-regulated genes). As a result, the results of pathway enrichment analyses, using the lists obtained from the t-test analysis, were very similar to those using the lists obtained from the multiple linear regression analysis.

#### *Correlation analysis between SMRI differential genes' expression and subjects' information regarding medications, substance and alcohol use*

Correlation analyses between the expression of the SMRI 881 up-regulated genes and 986 down-regulated genes and Fluphenazine equivalent dosage, severity of substance use and severity of alcohol use was performed. The Pearson correlation histogram for the SMRI 986 down-regulated genes is plotted in Figures 2S-4S for these parameters. In addition, p-values for each correlation value was calculated. For both the down-regulated genes and up-regulated genes, and for each of the 3 parameters (Fluphenazine equivalent, substance use and alcohol use), FDR(6) was applied and corrected p-values were calculated. While for the 881 up-regulated genes no gene passed FDR of 5% in each of the 3 parameters, for the 986 down-regulated genes 2 genes passed FDR of 5% for substance use, ADSL and C9orf85.

#### *Gene expression meta-analysis*

For a given gene, a meta-analysis that integrates its expression in both SMRI and MSSM was applied. To address the differences in study design and platform usage, we applied the Effect size (ES), the standardized difference between the expression in the disease vs. control samples, combined with Random Effect Modeling, which takes both the direction and magnitude of gene expression changes into consideration to generate more biologically consistent results. As was demonstrated in (7), it is superior to other meta-analytic methods in that it has the ability to handle the variability between studies, and highly applicable for gene expression data. ES (Hedges' g (8)) was calculated separately for the SMRI and MSSM datasets. The direction of the effect size was positive if the expression in the disease group was higher than in the control group. Hedges'g and confidence interval values were calculated for each of the SMRI and MSSM datasets, using the function "metacont" from the "meta" package in R, a general package for meta-analysis, version 4.9-2 (9). The summary measure of the two datasets with its confidence interval was calculated using the same function, using the random effects model (10).

As an example, results of the meta-analysis of proteasome subunit α5, PSMA5, is plotted in Figure 6S. It can be seen that while in each of the SMRI and MSSM separately we can observe only a trend towards down-regulation (95% confidence interval horizontal lines cross the zero), statistical significance was achieved only when the two datasets were integrated.

### *Additional datasets characteristics*

#### SMRI dataset

**Brodmann Area 23 (BA23) SMRI samples:** BA23 postmortem tissues from 13 subjects with schizophrenia and 15 healthy controls were obtained from the SMRI using approved protocols for tissue collection and informed consent (11). All samples were examined by a certified neuropathologist to exclude Alzheimer's disease and other cerebral pathology (11). Diagnoses were performed independently by two psychiatrists according to DSM-IV criteria. See samples' characteristics in Table 2S.

**BA23 SMRI RNA extraction and quality control:**The brain regions were dissected and total RNA was isolated using the Trizol method by the staff at SMRI. The concentration of total RNA and RNA Integrity Number value (RIN) were measured. Total RNA samples were delivered on dry ice to The Nancy & Stephen Grand Israel National Center for Personalized Medicine (G-INCPM) for whole transcriptome sequencing. Samples with RIN ⩾5 were selected for sequencing (all 28 samples). Among these samples, the mean RIN was 8.2 (± 0.5).

**STG and BA23 SMRI RNA sequencing libraries preparation:** Libraries preparation were done using the INCPM-RNA-seq protocol. Briefly, polyA fraction (mRNA) was purified from 500ng of total RNA by oligo(dT) beads following by fragmentation and generation of double stranded cDNA using random hexamers. Then, end repair, A base addition, adapter ligation and PCR amplification steps were performed. Libraries were evaluated by Qubit and TapeStation and pooled in an equimolar ratio. Sequencing libraries were constructed with barcodes to allow multiplexing of samples in a lane. For raw RNA sequencing data description see Table 1S.

**STG and BA23 SMRI mapping and quantification of gene expression:** Fragment mapped to the genome (hg19) using TopHat version V2.0.5 and Bowtie version 2.2.0. Only fragments with good quality reads (mean Qphred per read>20; corresponding to above 99% probability of a correctly identified base), with both ends uniquely mapped to the genome, were considered (~70% of all fragments). The signals from the 6 lanes were summed. Known Ensembl gene levels were quantified by HTSeq version 0.6.0 in intersection-strict mode. This provides an integral count of reads for each gene in each sample (a sample-by-gene ‘read count matrix’). Gene models were downloaded from the UCSC Genome Browser (http://genome.ucsc.edu/; assembly hg19).

**BA23 SMRI preprocessing:** Lowess correction was calculated (12). Then expression threshold was set to 6 (log scale) to reduce noise. Filtering: Genes with expression values below 6 in at least 80% of the samples (considering both STG and BA23 SMRI samples) were filtered out of the analysis. 16,482 genes were left for the rest of the analysis after filtering (out of 23,715).

**Table 1S. SMRI STG and BA23 RNA-seq data**

| Group | N | Mean total reads/subject | Mapped reads (%) | No. of genes sequenced |
|---|---|---|---|---|
| **STG, SMRI dataset** | | | | |
| Schizophrenia | 14 | 23,559,907 | 91.2 | 15,554 |
| Control | 15 | 22,976,936 | 92.1 | 15,635 |
| **BA23, SMRI dataset** | | | | |
| Schizophrenia | 13 | 29,004,007 | 92.3 | 15,818 |
| Control | 15 | 41,158,436 | 92.4 | 15,804 |

### STG, Barnes 2011 dataset

The dataset GSE21935 (13) was downloaded from the GEO database (https://www.ncbi.nlm.nih.gov/geo/query/acc.cgi?acc=GSE21935). The dataset consists of 42 superior temporal cortex samples from subjects with schizophrenia (n=23) and healthy controls (n=19). Samples were run on Affymetrix Human Genome U133 Plus 2.0 Arrays. See Table 2S for samples’ characteristics. **Normalization method:** Arrays were scanned on a GeneChip Scanner 3000, and fluorescence intensity was obtained by using GeneChip Operating Software (13). As described in GSE21935_series_matrix.txt (available at https://www.ncbi.nlm.nih.gov/geo/query/acc.cgi?acc=GSE21935), the data were analyzed with Microarray Suite version 5.0 (MAS 5.0) using Affymetrix default analysis settings and global scaling as normalization method. The trimmed mean target intensity of each array was arbitrarily set to 150. We then applied threshold and logarithm base 2 (log2). The threshold value was determined using scatter plots of healthy control samples, in order to estimate the noise level (the threshold after log2 that was used is 4). **Filtering:** 1) Initial number of probe-sets was 54,675 (45,772 with assigned gene symbols). 2) In case of gene symbol with multiple probe-sets, the probe-set with the highest mean expression over the samples was taken into account and the other probe-sets were discarded. Number of genes after this step: 22,880. 3) Genes that are absent (values equal or lower than the threshold) in more than 70% of both the schizophrenia and the control samples, were filtered out. Number of genes after filtering: 17,464.

#### Cerebellum, Chen 2018 dataset

The dataset GSE35978 (14) was downloaded from the GEO database (https://www.ncbi.nlm.nih.gov/geo/query/acc.cgi?acc=GSE35978). The initial dataset consisted of 312 brain samples. We used only the cerebellum samples of subjects with schizophrenia (n=44) and healthy controls (n=50). See Table 2S for samples' characteristics. Samples were run on Affymetrix Human Gene 1.0 ST Array [transcript (gene) version]. **Normalization method:** As described in GSE35978_series_matrix.txt (available at https://www.ncbi.nlm.nih.gov/geo/query/acc.cgi?acc=GSE35978), the data were analyzed by Robust Multi-array Average (RMA) (15) using Affymetrix Expression Console with default analysis settings. **Filtering:** 1) Initial number of probe-sets was 33,297 (25,293 with assigned gene symbols). 2) In case of gene symbol with multiple probe-sets, the probe-set with the highest mean expression over the samples was taken into account and the other probe-sets were discarded. Number of genes after this step: 23,307. No threshold was applied, as the RMA algorithm in includes background correction and quantile normalization (16). Number of genes after filtering: 23,307 (no probes were removed).

#### DLPFC, Arion 2015 dataset

The dataset GSE93987 (15) was downloaded from the GEO database (https://www.ncbi.nlm.nih.gov/geo/query/acc.cgi?acc=GSE93987). The dataset consists of 208 dorsolateral prefrontal cortex (DLPFC) brain samples from subjects with schizophrenia (n=102) and healthy controls (n=106). See Table 2S for samples' characteristics. Samples were run on Affymetrix HT HG-U133 Arrays + PM Array Plate. **Normalization method:** Affymetrix CEL files were normalized and log2 transformed using RMA (15). **Filtering:** 1) Initial number of probe-sets: 54,613 (44,228 with assigned gene symbols). 2) In case of a gene symbol with multiple probe-sets, the probe-set with the highest mean expression over the samples was taken into account and the other probe-sets were discarded. Number of genes after this step: 21,597. No threshold was applied, as the RMA algorithm in includes background correction and quantile normalization (16). Number of genes after filtering: 21, 597 (no probes were removed).

### DLPFC, Ramaker 2017 dataset

The dataset GSE80655 (17) was downloaded from the GEO database (https://www.ncbi.nlm.nih.gov/geo/query/acc.cgi?acc=GSE80655). The initial dataset consisted of 281 brain samples. We used only the DLPFC of subjects with schizophrenia (n=24) and healthy controls (n=24). See Table 2S for samples' characteristics. Samples were run on Illumina HiSeq 2000 Arrays. **Normalization method:** To quantify the expression of each gene, RNA-seq reads were processed with aRNApipe v1.1 using default settings (17). We then applied threshold and log2. The threshold value was determined using scatter plots of control samples in order to estimate the noise level (the threshold after log2 that was used is 3). **Filtering:** 1) Initial number of probe-sets was 57,905 (21,287 with assigned gene symbols). 2) In case of a gene symbol with multiple probe-sets, the probe with the highest mean expression over the samples was taken into account and the other probe-sets were discarded. Number of genes after this step: 20,881. 3) Genes that are absent (values equal or lower than the threshold) in more than 70% of both the schizophrenia and the control samples, were filtered out. Number of genes after filtering: 17,037.

### BA10, Mycox 2009 dataset

The dataset GDS4523 (18) was downloaded from the GEO database (https://www.ncbi.nlm.nih.gov/sites/GDSbrowser?acc=GDS4523). The dataset consists of 51 BA10 brain samples from subjects with schizophrenia (n=28) and healthy controls (n=23). See Table 2S for samples' characteristics. Samples were run on Affymetrix Human Genome U133 Plus 2.0 Arrays. **Normalization method:** Arrays were scanned on a GeneChip Scanner 3000 and fluorescence intensity for each feature of the array was obtained by using GeneChip Operating Software (Affymetrix) (18). As described in GSE17612_series_matrix.txt (available at https://www.ncbi.nlm.nih.gov/sites/GDSbrowser?acc=GDS4523), the data were analyzed with Microarray Suite version 5.0 (MAS 5.0) using Affymetrix default analysis settings and global scaling as normalization method. The trimmed mean target intensity of each array was arbitrarily set to 150. We then applied threshold and log2. The threshold value was determined using scatter plots of healthy control samples in order to estimate the noise level (the threshold after log2 that was used is 4). **Filtering:** 1) The initial number of probe-sets was 54,613. 2) In case of a gene symbol with multiple probe-sets, the probe-set with the highest mean expression over the samples was taken into account and the other probe-sets were discarded. Number of genes after this step: 30,805. 3) Genes that are absent (values equal or lower than the threshold) in more than 70% of both the schizophrenia and the control samples, were filtered out. Number of genes after filtering: 27,295.

**Table 2S: Additional datasets characteristics; Average values ± standard deviation**

| Characteristics | Control | Schizophrenia | P-value |
|---|---|---|---|
| **BA23, SMRI dataset** | | | |
| Number of subjects | 15 | 13 | |
| Age (years) | 48.06 ±10.6 | 43.2 ±13.5 | 0.29 |
| Gender | 9M : 6F | 9M : 4F | 0.62 |
| Brain pH | 6.26 ±0.24 | 6.17 ± 0.26 | 0.36 |
| PMI | 23.7 ±9.9 | 33.9 ± 15.6 | 0.05 |
| **STG, Barnes 2011 dataset** (13), **GSE21935** | | | |
| Number of subjects | 19 | 23 | |
| Age (years) | 67.68 ± 22 | 72.17 ± 17 | 0.46 |
| Gender | 10M : 9F | 13M : 10F | 0.81 |
| Brain pH | 6.489 ± 0.32 | 6.161 ± 0.17 | 0.00013 |
| PMI | 9.105 ± 4.3 | 7.13 ± 5.7 | 0.22 |
| **Cerebellum, Chen 2018 dataset** (14), **GSE35978** | | | |
| Number of subjects | 50 | 44 | |
| Age (years) | 45.8 ± 9.3 | 43.18 ± 9.5 | 0.18 |
| Gender | 31M : 19F | 32M : 12F | 0.27 |
| Brain pH | 6.474 ± 0.32 | 6.428 ± 0.25 | 0.44 |
| PMI | 27.58 ± 11 | 33.27 ± 15 | 0.042 |
| **DLPFC, Arion 2015 dataset** (15), **GSE93987** | | | |
| Number of subjects | 106 | 102 | |
| Age (years) | Not provided | Not provided | |
| Gender | Not provided | Not provided | |
| Brain pH | Not provided | Not provided | |
| PMI | Not provided | Not provided | |
| **DLPFC, Ramaker 2017 dataset** (17), **GSE80655** | | | |
| Number of subjects | 24 | 24 | |
| Age (years) | 50.25 ± 13 | 42.67 ± 10 | 0.025 |
| Gender | 21M : 3F | 21M : 3F | 1 |
| Brain pH | 6.921 ± 0.11 | 6.83 ± 0.18 | 0.043 |
| PMI | 21.89 ± 6.6 | 21.29 ± 9.2 | 0.8 |
| **BA10, Mycox 2009 dataset** (18), **GDS4523** | | | |
| Number of subjects | 23 | 28 | |
| Age (years) | 69.04 ±22 | 73.32 ±15 | 0.41 |
| Gender | 12/11 | 19/9 | 0.26 |
| Brain pH | 6.15 ±0.29 | 6.15 ±0.21 | 8e-06 |
| PMI | 9.902 ± 4.4 | 8.714 ± 7 | 0.48 |
| **Overall number of subjects** | 237 | 234 | |

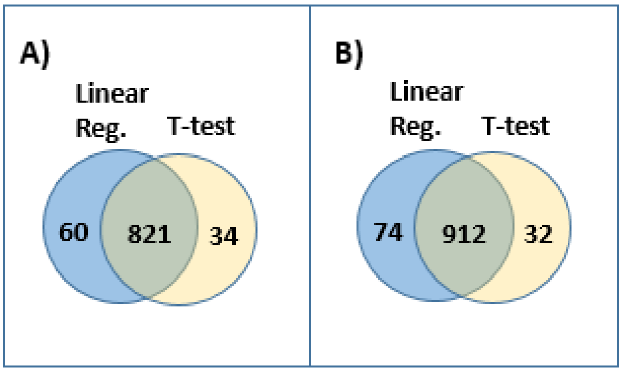

**Figure 1S: Comparison between multiple linear regression and t-test analysis resulting SMRI STG up- regulated and down-regulated genes. A)** Venn diagram for the intersection between the 881 genes that were found to be up-regulated in schizophrenia STG SMRI samples using multiple linear regression analysis (with FDR Q < 15%) and 855 genes that were found to be up-regulated in schizophrenia STG SMRI samples using t-test analysis (FDR Q < 15%). **B)** Venn diagram for the intersection between the 986 genes that were found to be down-regulated in schizophrenia STG SMRI samples using multiple linear regression analysis (with FDR Q < 15%) and 944 genes that were found to be down-regulated in schizophrenia STG SMRI samples using t-test analysis (FDR Q < 15%).

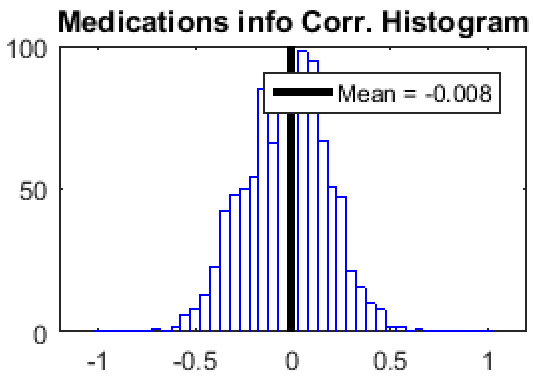

**Figure 2S: SMRI STG 986 Down-regulated genes: Pearson Correlation Histogram between lifetime quantity of Fluphenazine or equivalent antipsychotic (in mg) and gene expression, along the 14 schizophrenia patients for which this information is available.** The X-axis represents the Pearson correlation values, the mean correlation value measured for the 986 down-regulated genes is specified by a black vertical line.

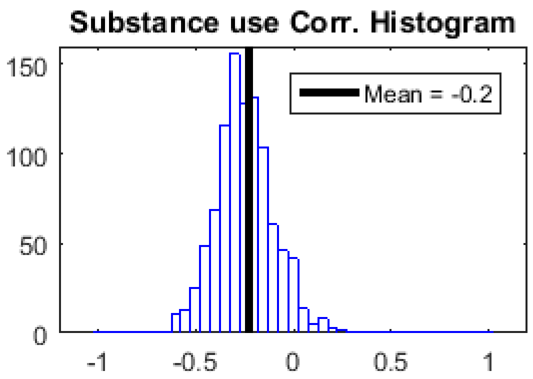

**Figure 3S: SMRI STG 986 Down-regulated genes: Pearson Correlation Histogram between Substance use severity (measured 0-5) and gene expression, measured along schizophrenia and control subjects.** The X-axis represents the Pearson correlation values, the mean correlation value measured for the 986 down-regulated genes is specified by a black vertical line.

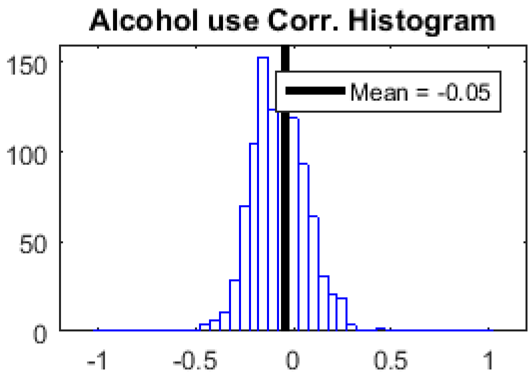

**Figure 4S: SMRI STG 986 Down-regulated genes: Pearson Correlation Histogram between Substance use severity (measured 0-5) and gene expression, measured along schizophrenia and control subjects.** The X-axis represents the Pearson correlation values, the mean correlation value measured for the 986 down-regulated genes is specified by a black vertical line.

A)

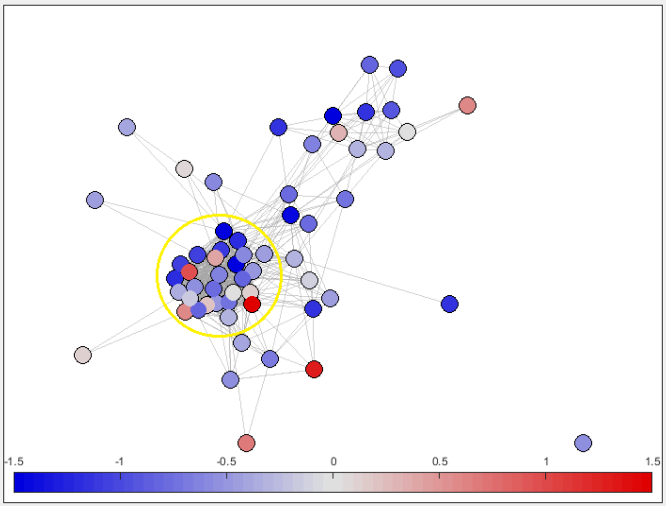

B)

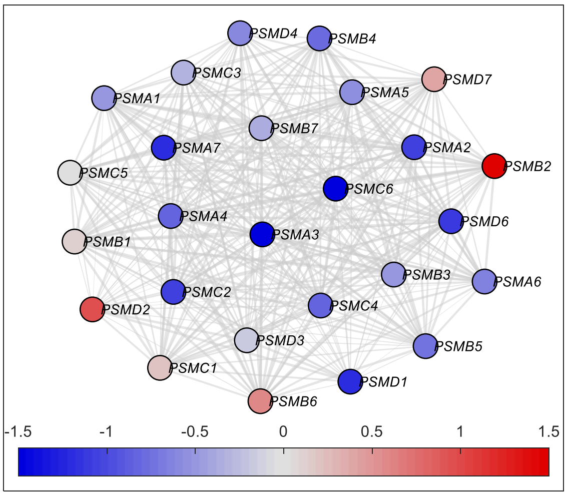

**Figure 5S: MSSM STG Differential expression network view: A) Ubiquitin-Proteasome Dependent Proteolysis superPathway.** The node's colors correspond to the deviation from the control samples group, in terms of standard deviation units (see Methods). The edges represent STRING database co-expression relations. Only genes that have co-expression relations with other genes in the network are displayed. A subgroup of highly-interconnected genes, coding for proteasome subunits, is circled. **B) Zoom in on proteasome subunits.** The same plot as in A), for a subgroup of highly-interconnected genes coding for proteasome subunits (circled in A)).

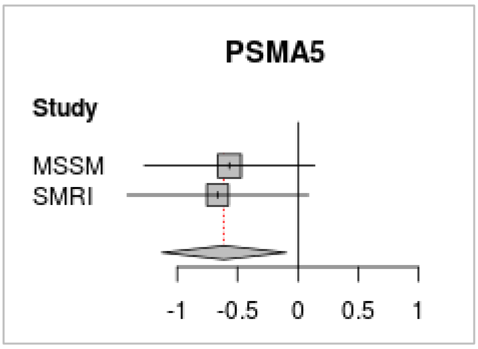

**Figure 6S: PSMA5 STG SMRI + MSSM meta-analysis of differential expression.** Forest plot was generated using the function “forest” from the “meta” package in R, version 4.9-2 (General Package for Meta-Analysis) (9). The forest plot shows the differences in PSMA5 expression between subjects with schizophrenia and healthy controls, for each of the two studies, SMRI and MSSM. Each square represents the standardized difference (Hedges’ g (8)) between schizophrenia and control for that study, with the area of the square reflecting the weight (determined by the sample size) given to that study in the meta-analysis. Each horizontal line represents the 95% confidence interval for the mean difference in that study. The vertical line shows the point of 0 difference. The standardized difference is positive (negative) if the expression is higher (lower) in schizophrenia vs. the control group. The center of the diamond represents the overall difference across both studies and its width represents 95% confidence interval.

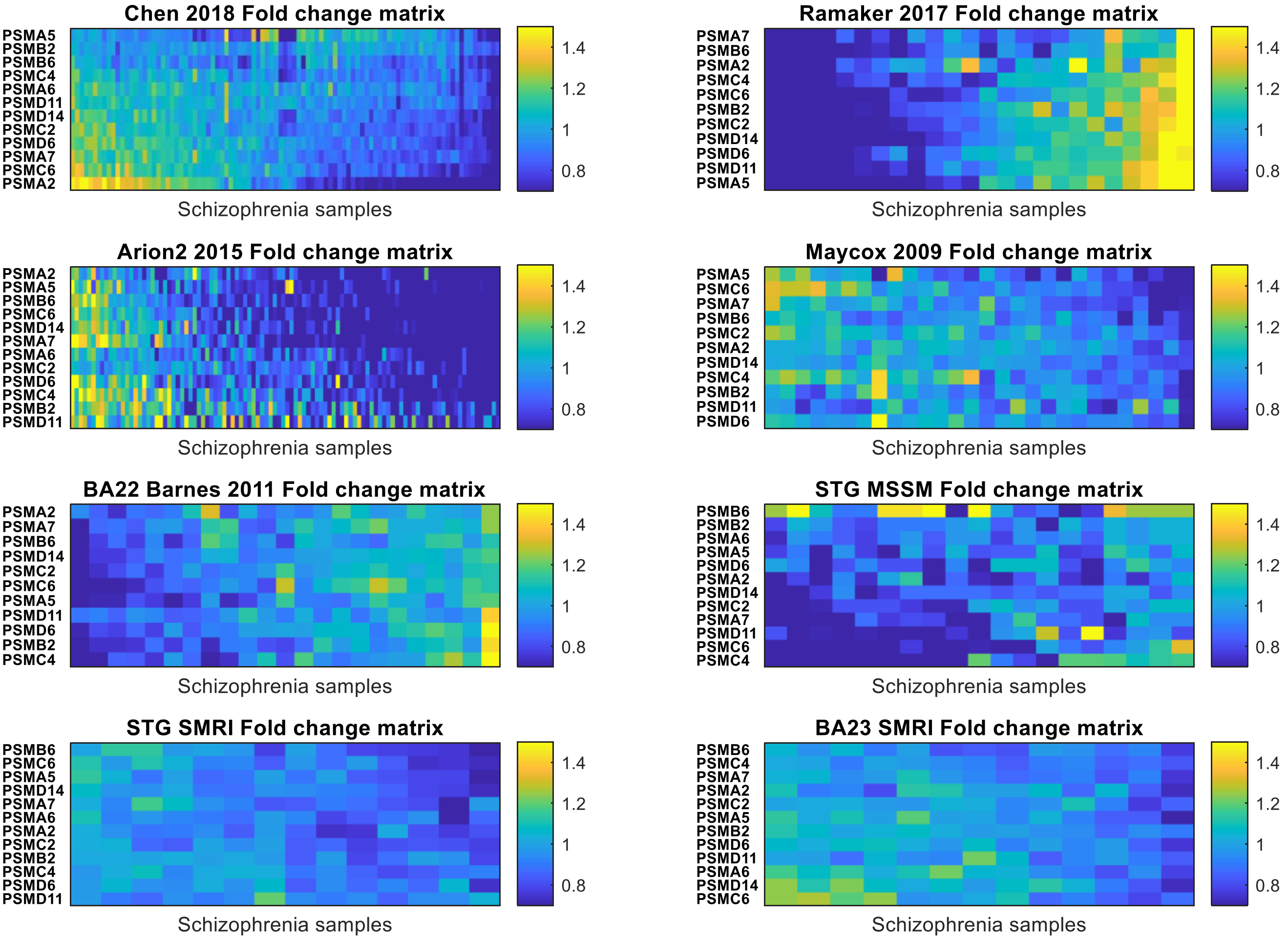

**Figure 7S: Fold change matrix of proteasome subunits genes. A) Cerebellum, Chen 2018 data** (14). Each row represents one of the 12 proteasome subunits genes that were found to be down-regulated in schizophrenia in the meta-analysis of the SMRI and MSSM datasets. Each column represents one of the Chen 2018 44 schizophrenia samples. The color code represents the fold change, i.e. the expression value of the proteasome subunit gene in the specific sample, divided by its mean expression in the 50 control samples. **B) DLPFC, Ramaker 2017 dataset** (17). 24 samples of schizophrenia patients vs. 24 controls. **C) DLPFC, Arion 2015 dataset** (15). 102 samples of schizophrenia patients vs. 106 controls. **D) BA10, Mycox 2009 dataset** (18). 28 samples of schizophrenia patients vs. 23 controls. **E) STG SMRI dataset.** 14 samples of schizophrenia patients vs. 15 controls. **F) BA23 SMRI dataset.** 13 samples of schizophrenia patients vs. 15 controls.

**Table 3S: Pathway enrichment analysis of SMRI STG up-regulated genes with corrected p-value < 0.05.** GeneAnalytics tool superpathways that were found to be enriched in the list of up-regulated genes are ordered by descending order of their enrichment score. The number of up-regulated genes that belong to each superpathway is given, with the number of genes of each superpathway in parentheses.

| # | Score | SuperPath Name | Num Matched (SuperPath) genes | MSSM Enrichment Score |
|---|---|---|---|---|
| 1 | 13.43 | Metallothioneins Bind Metals | 5 (11) | - |
| 2 | 13.2 | Axon Guidance | 19 (175) | - |
| 3 | 12.4 | Protein-protein Interactions at Synapses | 11 (72) | - |
| 4 | 11.42 | ERK Signaling | 71 (1177) | 18.14 |
| 5 | 10.38 | Influenza Viral RNA Transcription and Replication | 16 (158) | 18.85 |
| 6 | 10.29 | NFAT and Cardiac Hypertrophy | 26 (326) | - |
| 7 | 10.25 | Hedgehog Pathway | 10 (73) | - |
| 8 | 10.2 | 4-hydroxytamoxifen, Dexamethasone, and Retinoic Acids Regulation of P27 Expression | 5 (18) | - |
| 9 | 10.14 | P38 MAPK Signaling Pathway (sino) | 9 (61) | - |
| 10 | 9.92 | RET Signaling | 59 (974) | - |
| 11 | 9.4 | CREB Pathway | 36 (528) | 13.49 |
| 12 | 9.25 | LKB1 Signaling Events | 7 (42) | - |
| 13 | 9.01 | Phospholipase-C Pathway | 34 (498) | 10.15 |
| 14 | 8.8 | Signaling By NOTCH1 | 12 (113) | - |
| 15 | 8.76 | Circadian Entrainment | 21 (262) | - |
| 16 | 8.74 | MAPK Signaling Pathway | 24 (316) | - |
| 17 | 8.74 | HIV Life Cycle | 52 (865) | 12.88 |
| 18 | 8.7 | Regulation of Lipid Metabolism Insulin Signaling-generic Cascades | 21 (263) | - |
| 19 | 8.57 | P38 MAPK Signaling Pathway (WikiPathways) | 6 (34) | - |
| 20 | 8.56 | Focal Adhesion | 22 (283) | 11.03 |

**Table 4S. Pathway enrichment analysis of SMRI STG down-regulated genes with corrected p-value < 0.05.** GeneAnalytics tool superpathways that were found to be enriched in the list of down-regulated genes are ordered by descending order of their enrichment score. The enrichment scores are in the second column and the superpathways' names are listed in the third column. The fourth column presents the number of down-regulated genes that belong to each superpathway , with the total number of genes of the superpathway in parentheses. MSSM enrichment score is given in the 5th column, where (-) sign means that the superpathway wasn't enriched in the list of MSSM down-regulated genes. For superpathways that are known to involve the UPS, a reference indicating the UPS involvement is given in the 6th column. Ubiquitin-proteasome directly related pathways are in bold

| # | Score | SuperPath Name | Num Matched (SuperPath) genes | MSSM Enrichment Score | Evidence for UPS involvement |
|---|---|---|---|---|---|
| 1 | 41.07 | MRNA Splicing - Major Pathway | 65 (307) | 33.59 | |
| 2 | 28.4 | Chks in Checkpoint Regulation | 46 (224) | 18.43 | |
| 3 | 27.72 | Translational Control | 41 (189) | 24.61 | |
| 4 | 26.05 | Vesicle-mediated Transport | 93 (660) | 18.05 | |
| 5 | 25.02 | CDK-mediated Phosphorylation and Removal of Cdc6` | 114 (880) | 18.12 | the UPS plays a central role (19) |
| 6 | 24.58 | Gene Expression | 203 (1841) | 29.46 | |
| 7 | 21.52 | Protein Processing in Endoplasmic Reticulum | 34 (166) | 20.01 | integrally involved in the UPS (20) |
| 8 | 21.33 | DNA Damage | 49 (292) | 13.79 | closely involve the UPS (21) |
| 9 | 21.24 | Cell Cycle, Mitotic | 84 (622) | 13.65 | tightly regulated by the UPS (22) |
| **10** | **19.58** | **Ubiquitin-Proteasome Dependent Proteolysis** | **27 (122)** | **18.4** | |
| **11** | **19.06** | **Metabolism of Proteins** | **175 (1628)** | **25.5** | |
| 12 | 18.59 | Regulation of Degradation of DeltaF508 CFTR in CF | 18 (63) | 10.93 | dominated by the UPS (23) |
| 13 | 16.6 | Cell Cycle | 28 (145) | - | |
| **14** | **16.59** | **Ubiquitin Mediated Proteolysis** | **27 (137)** | **10.94** | |
| 15 | 16.23 | Signaling By Hedgehog | 27 (139) | 12.4 | |
| **16** | **15.57** | **Proteolysis_Putative Ubiquitin Pathway** | **12 (35)** | **-** | |
| 17 | 15.39 | Cellular Response to Heat Stress | 20 (89) | 13.16 | heat shock proteins recognize misfolded proteins and incorporate the UPS (24) |
| 18 | 15.35 | Transcription-Coupled Nucleotide Excision Repair (TC-NER) | 23 (112) | - | |
| 19 | 15.29 | Nucleotide Excision Repair | 16 (61) | - | |
| 20 | 14.6 | Class I MHC Mediated Antigen Processing and Presentation | 95 (823) | 15.89 | |
| 21 | 14.52 | Innate Immune System | 210 (2132) | 22.32 | |
| 22 | 14.01 | HIV Life Cycle | 98 (865) | 11.41 | |
| 23 | 13.78 | Clathrin-mediated Endocytosis | 25 (137) | - | a key process that transports a wide |

| | | | | | |
|---|---|---|---|---|---|
| | | | | | range of molecules from the cell surface to the interior and is closely regulated by the UPS (25) |
| 24 | 13.68 | Signaling By NOTCH1 | 22 (113) | - | |
| 25 | 13.47 | Copper Homeostasis | 14 (54) | - | |
| 26 | 13.46 | Mitotic G1-G1/S Phases | 27 (156) | - | |
| 27 | 13.29 | Transport to The Golgi and Subsequent Modification | 41 (285) | 17.23 | |
| 28 | 13.2 | Regulation of Cholesterol Biosynthesis By SREBP (SREBF) | 14 (55) | - | |
| 29 | 13.13 | Telomere C-strand (Lagging Strand) Synthesis | 20 (100) | - | |
| 30 | 13 | Terpenoid Backbone Biosynthesis | 11 (36) | - | |
| 31 | 12.61 | Processing of Capped Intronless Pre-mRNA | 10 (31) | - | |
| 32 | 12.56 | Mitotic Metaphase and Anaphase | 29 (180) | 11.56 | tightly regulated by the UPS (26) |
| 33 | 12.45 | Transport of The SLBP Independent Mature MRNA | 31 (199) | 12.88 | |
| 34 | 12.31 | Remodeling of Adherens Junctions | 22 (121) | 12.71 | cadherin, the main adhesion molecule in adherens junctions, is tightly regulated by the UPS (27) |
| 35 | 12.08 | Cell Cycle Checkpoints | 31 (202) | - | |
| 36 | 11.92 | Presenilin Action in Notch and Wnt Signaling | 12 (46) | - | |
| 37 | 11.5 | Circadian Rythm Related Genes | 31 (207) | 13.68 | |
| 38 | 11.4 | RNA Transport | 27 (171) | 14.03 | |
| 39 | 11.3 | CLEC7A (Dectin-1) Signaling | 24 (145) | 11.77 | |
| 40 | 11.17 | Formation of HIV Elongation Complex in The Absence of HIV Tat | 28 (182) | - | |
| 41 | 10.96 | Cellular Senescence | 55 (452) | 14.27 | |
| 42 | 10.95 | RNA Polymerase II Transcription Termination | 15 (72) | - | |
| 43 | 10.94 | Calnexin/calreticulin Cycle | 10 (36) | - | |
| 44 | 10.86 | Mechanisms of CFTR Activation By S-nitrosoglutathione (normal and CF) | 11 (43) | 9.96 | |
| **45** | **10.75** | **Proteolysis Role of Parkin in The Ubiquitin-Proteasomal Pathway** | **15 (73)** | **15.85** | |
| 46 | 10.75 | Sterol Regulatory Element-Binding Proteins (SREBP) Signalling | 15 (73) | - | |
| 47 | 10.67 | Metabolism | 235 (2543) | 10.35 | |
| 48 | 10.52 | Cytoskeletal Signaling | 40 (304) | 12.9 | |
| 49 | 10.24 | Brain-Derived Neurotrophic Factor (BDNF) Signaling Pathway | 23 (144) | 11.87 | |

**Table 5S: Pathway enrichment analysis of MSSM STG up-regulated genes with corrected p-value < 0.05.** GeneAnalytics tool superpathways that were found to be enriched in the list of up-regulated genes are ordered by descending order of their enrichment score. The number of up-regulated genes that belong to each superpathway is given, with the number of genes of each superpathway in parentheses.

| # | Score | SuperPath Name | Num Matched (SuperPath) genes |
|---|---|---|---|
| 1 | 23.36 | GPCR Pathway | 59 (708) |
| 2 | 20.4 | Metabolism | 148 (2544) |
| 3 | 18.85 | Influenza Viral RNA Transcription and Replication | 21 (158) |
| 4 | 18.14 | ERK Signaling | 79 (1177) |
| 5 | 17.08 | Metabolism of Proteins | 100 (1628) |
| 6 | 17.07 | PEDF Induced Signaling | 54 (721) |
| 7 | 16.36 | Degradation of The Extracellular Matrix | 29 (298) |
| 8 | 15.13 | RRNA Processing in The Nucleus and Cytosol | 22 (203) |
| 9 | 14.93 | Influenza A | 29 (315) |
| 10 | 14.54 | Pathways in Cancer | 41 (528) |
| 11 | 13.8 | TGF-Beta Pathway | 47 (652) |
| 12 | 13.49 | CREB Pathway | 40 (528) |
| 13 | 13.31 | Integrin Pathway | 42 (568) |
| 14 | 12.88 | HIV Life Cycle | 57 (865) |
| 15 | 12.2 | Regulation of Insulin-like Growth Factor (IGF) Transport and Uptake By Insulin-like Growth Factor Binding Proteins (IGFBPs) | 6 (21) |
| 16 | 11.18 | Naphthalene Metabolism | 6 (24) |
| 17 | 11.06 | Neuropathic Pain-Signaling in Dorsal Horn Neurons | 21 (232) |
| 18 | 11.03 | Focal Adhesion | 24 (283) |
| 19 | 10.66 | Actin Nucleation By ARP-WASP Complex | 27 (341) |
| 20 | 10.64 | Akt Signaling | 45 (681) |
| 21 | 10.6 | PAK Pathway | 45 (682) |
| 22 | 10.46 | Apoptotic Pathways in Synovial Fibroblasts | 47 (725) |
| 23 | 10.37 | FOXM1 Transcription Factor Network | 7 (37) |
| 24 | 10.29 | NRF2 Pathway | 15 (145) |
| 25 | 10.24 | Cell Adhesion_ECM Remodeling | 9 (61) |
| 26 | 10.18 | G-Beta Gamma Signaling | 27 (349) |
| 27 | 10.15 | Phospholipase-C Pathway | 35 (498) |

**Table 6S: Pathway enrichment analysis of MSSM STG down-regulated genes with corrected p-value < 0.05.** GeneAnalytics tool superpathways that were found to be enriched in the list of down-regulated genes are ordered by descending order of their enrichment score. The number of down-regulated genes that belong to each superpathway is given, with the number of genes of each superpathway in parentheses. UPS directly related pathways are in bold

| # | Score | SuperPath Name | Num Matched (SuperPath) genes |
|---|---|---|---|
| 1 | 33.59 | MRNA Splicing - Major Pathway | 38 (307) |
| 2 | 29.46 | Gene Expression | 116 (1841) |
| **3** | **25.5** | **Metabolism of Proteins** | **102 (1628)** |
| 4 | 24.61 | Translational Control | 25 (189) |
| 5 | 22.32 | Innate Immune System | 121 (2132) |
| 6 | 20.01 | Protein Processing in Endoplasmic Reticulum | 21 (166) |
| 7 | 18.43 | Chks in Checkpoint Regulation | 24 (224) |
| **8** | **18.4** | **Ubiquitin-Proteasome Dependent Proteolysis** | **17 (122)** |
| 9 | 18.12 | CDK-mediated Phosphorylation and Removal of Cdc6 | 59 (880) |
| 10 | 18.05 | Vesicle-mediated Transport | 48 (660) |
| 11 | 17.23 | Transport to The Golgi and Subsequent Modification | 27 (285) |
| 12 | 15.89 | Class I MHC Mediated Antigen Processing and Presentation | 54 (823) |
| **13** | **15.85** | **Proteolysis Role of Parkin in The Ubiquitin-Proteasomal Pathway** | **12 (73)** |
| 14 | 14.27 | Cellular Senescence | 34 (452) |
| 15 | 14.03 | RNA Transport | 18 (171) |
| 16 | 13.93 | Beta-Adrenergic Signaling | 26 (308) |
| 17 | 13.79 | DNA Damage | 25 (292) |
| 18 | 13.75 | Telomere Extension By Telomerase | 6 (19) |
| 19 | 13.68 | Circadian Rythm Related Genes | 20 (207) |
| 20 | 13.65 | Cell Cycle, Mitotic | 42 (622) |
| 21 | 13.16 | Cellular Response to Heat Stress | 12 (89) |
| 22 | 12.9 | Cytoskeletal Signaling | 25 (304) |
| 23 | 12.88 | Transport of The SLBP Independent Mature MRNA | 19 (199) |
| 24 | 12.71 | Remodeling of Adherens Junctions | 14 (121) |
| 25 | 12.49 | Integrated Breast Cancer Pathway | 16 (154) |
| 26 | 12.4 | Signaling By Hedgehog | 15 (139) |
| 27 | 12.19 | Immune Response_IL-6 Signaling Pathway | 7 (33) |
| 28 | 12.1 | Ran Pathway | 5 (15) |
| 29 | 11.87 | Brain-Derived Neurotrophic Factor (BDNF) Signaling Pathway | 15 (144) |
| 30 | 11.86 | Signaling By Wnt | 26 (338) |
| 31 | 11.77 | CLEC7A (Dectin-1) Signaling | 15 (145) |
| 32 | 11.67 | RNA Polymerase II Transcription Initiation And Promoter Clearance | 19 (213) |
| 33 | 11.56 | Mitotic Metaphase and Anaphase | 17 (180) |
| 34 | 11.41 | HIV Life Cycle | 51 (865) |
| 35 | 11.39 | ErbB1 Downstream Signaling | 12 (102) |
| 36 | 11.3 | Viral MRNA Translation | 36 (547) |
| 37 | 11.21 | Cytokine Signaling in Immune System | 46 (761) |

| | | | |
|---|---|---|---|
| 38 | 10.95 | Antigen Processing-Cross Presentation | 13 (121) |
| **39** | **10.94** | **Ubiquitin Mediated Proteolysis** | **14 (137)** |
| 40 | 10.93 | Regulation of Degradation of DeltaF508 CFTR in CF | 9 (63) |
| **41** | **10.59** | **Deubiquitination** | **23 (301)** |
| 42 | 10.56 | TRNA Processing | 12 (109) |
| 43 | 10.35 | Metabolism | 120 (2543) |
| 44 | 10.25 | Sulfur Amino Acid Metabolism | 8 (54) |
| 45 | 10.25 | Translation Factors | 8 (54) |
| 46 | 10.24 | WNT Signaling | 20 (250) |
| 47 | 10.07 | Apoptotic Pathways in Synovial Fibroblasts | 43 (725) |
| 48 | 9.96 | Mechanisms of CFTR Activation By S-nitrosoglutathione (normal and CF) | 7 (43) |

**Table 7S: STG down-regulated genes enriched UPS-related pathways; numbers of SMRI and MSSM hits.** For each of the 5 UPS-related pathways that were enriched in the SMRI STG down-regulated genes, the number of 'hits' (down-regulated genes that belong to the pathway) shared with MSSM STG down-regulated genes is given in the second column. The numbers of hits specific to SMRI and MSSM down-regulated genes are given in columns 3 and 4, respectively.

| UPS pathway | # SMRI+MSSM hits | # SMRI only hits | # MSSM only hits |
|---|---|---|---|
| Ubiquitin-Proteasome Dependent Proteolysis | 4 | 27 | 17 |
| Metabolism of Proteins | 27 | 235 | 120 |
| Ubiquitin Mediated Proteolysis | 5 | 27 | 14 |
| Proteolysis Role of Parkin in The Ubiquitin-Proteasomal Pathway | 2 | 15 | 12 |
| Proteolysis_Putative Ubiquitin Pathway | 0 | 12 | 0 (pathway wasn't enriched in MSSM) |

**Table 8S: SMRI STG Differential expression analysis of proteasome subunits genes. 7 schizophrenia samples ("Group 2") were compared to the 15 control samples.** T-sample t-test was applied; p-values were corrected by the benjamini-Hochberg procedure (6)

| # | Gene Symbol | t-statistic | p-value | Corrected p-value |
|---|---|---|---|---|
| 1 | PSMC4 | -5.08 | 5.75E-05 | 0.00282 |
| 2 | PSMA2 | -4.51 | 0.000215 | 0.00527 |
| 3 | PSMC2 | -4.28 | 0.000366 | 0.00597 |
| 4 | PSMB5 | -3.77 | 0.00121 | 0.0123 |
| 5 | PSMB6 | -3.75 | 0.00125 | 0.0123 |
| 6 | PSMD2 | -3.45 | 0.00253 | 0.0174 |
| 7 | PSMB2 | -3.42 | 0.00268 | 0.0174 |
| 8 | PSMD3 | -3.4 | 0.00285 | 0.0174 |
| 9 | PSMC6 | -3.35 | 0.0032 | 0.0174 |
| 10 | PSMD8 | -3.27 | 0.00386 | 0.0189 |
| 11 | PSME4 | -3.11 | 0.0055 | 0.0241 |
| 12 | PSMA7 | -3.08 | 0.0059 | 0.0241 |

| | | | | |
|---|---|---|---|---|
| 13 | PSMD14 | -2.84 | 0.0101 | 0.0381 |
| 14 | PSMB7 | -2.79 | 0.0112 | 0.0393 |
| 15 | PSMA5 | -2.74 | 0.0127 | 0.0394 |
| 16 | PSMA1 | -2.73 | 0.0129 | 0.0394 |
| 17 | PSMD13 | -2.48 | 0.0221 | 0.0636 |
| 18 | PSMA6 | -2.39 | 0.0266 | 0.0723 |
| 19 | PSME3 | -2.34 | 0.0298 | 0.0768 |
| 20 | PSMD6 | -2.21 | 0.0393 | 0.0963 |
| 21 | PSME2 | 2.17 | 0.0426 | 0.0995 |
| 22 | PSMD11 | -2.1 | 0.0484 | 0.107 |
| 23 | PSMB10 | 1.74 | 0.0975 | 0.199 |
| 24 | PSMG2 | 1.7 | 0.104 | 0.203 |
| 25 | PSMB9 | 1.57 | 0.132 | 0.25 |
| 26 | PSMB8 | 1.51 | 0.148 | 0.268 |
| 27 | PSMC1 | -1.48 | 0.155 | 0.271 |
| 28 | PSMG1 | -1.39 | 0.179 | 0.298 |
| 29 | PSMB1 | -1.38 | 0.182 | 0.298 |
| 30 | PSMA4 | -1.33 | 0.199 | 0.315 |
| 31 | PSMD7 | -1.23 | 0.234 | 0.349 |
| 32 | PSMC5 | -1.22 | 0.235 | 0.349 |
| 33 | PSMD5 | 1.14 | 0.27 | 0.389 |
| 34 | PSMA3 | -1.11 | 0.279 | 0.39 |
| 35 | PSMG3 | -0.95 | 0.353 | 0.481 |
| 36 | PSMG4 | -0.835 | 0.413 | 0.548 |
| 37 | PSMD9 | -0.75 | 0.462 | 0.595 |
| 38 | PSMB4 | -0.71 | 0.486 | 0.61 |
| 39 | PSMD1 | 0.645 | 0.526 | 0.645 |
| 40 | PSMD12 | -0.537 | 0.597 | 0.701 |
| 41 | PSMD4 | -0.532 | 0.601 | 0.701 |
| 42 | PSMD10 | 0.486 | 0.632 | 0.708 |
| 43 | PSMB3 | -0.481 | 0.636 | 0.708 |
| 44 | PSMF1 | -0.458 | 0.652 | 0.71 |
| 45 | PSME1 | 0.258 | 0.799 | 0.852 |
| 46 | PSMC3 | 0.157 | 0.877 | 0.914 |
| 47 | PSMC3IP | 0.0853 | 0.933 | 0.936 |
